# Seasonal and long-term consequences of esca on grapevine stem xylem integrity

**DOI:** 10.1101/2020.09.07.282582

**Authors:** G. Bortolami, E. Farolfi, E. Badel, R. Burlett, H. Cochard, N. Ferrer, A. King, L.J. Lamarque, P. Lecomte, M. Marchesseau-Marchal, J. Pouzoulet, J.M. Torres-Ruiz, S. Trueba, S. Delzon, G.A. Gambetta, C.E.L. Delmas

## Abstract

Hydraulic failure has been extensively studied during drought-induced plant dieback, but its role in plant-pathogen interactions is under debate. During esca, a grapevine (*Vitis vinifera*) disease, symptomatic leaves are prone to irreversible hydraulic dysfunctions but little is known about the hydraulic integrity of perennial organs over the short- and long-term. We investigated the effects of esca on stem hydraulic integrity in naturally infected plants within a single season and across season(s). We coupled direct (*k_s_*) and indirect (*k_th_*) hydraulic conductivity measurements, and tylose and vascular pathogen detection with in vivo X-ray microtomography visualizations. We found xylem occlusions (tyloses), and subsequent loss of stem *k_s_*, in all of the shoots with severe symptoms (apoplexy) and in more than 60% of the shoots with moderate symptoms (tiger-stripe), and no tyloses in shoots that were currently asymptomatic. In vivo stem observations demonstrated that tyloses were observed only when leaf symptoms appeared, and resulted in more than 50% PLC in 40% of symptomatic stems, unrelated to symptom age. The impact of esca on xylem integrity was only seasonal and no long-term impact of disease history was recorded. Our study demonstrated how and to what extent a vascular disease such as esca, affecting xylem integrity, could amplify plant mortality by hydraulic failure.

**Highlight:** Our study reveals that esca can critically affect xylem water movement in grapevine perennial organs, by the presence of plant-derived tyloses.

## INTRODUCTION

In agricultural and forest ecosystems, perennial plant dieback causes decreases in plant productivity and longevity (Aleemullah and Walsh, 1996; Eskalen *et al*., 2013; Urbez-Torres *et al*., 2013; Alvindia and Gallema, 2017). Plant dieback is a complex process where different biotic and/or abiotic stress factors interact and contribute to leaf and crown wilting and ultimately plant death (Desprez-Lostau *et al*., 2006; Anderegg *et al*., 2013, Cailleret *et al*., 2017; Bettenfeld *et al*., 2020). Drought-mediated plant dieback has been extensively studied, and in this case hydraulic failure has been identified as the primary cause of plant death (Anderegg *et al*., 2016). Hydraulic failure results from an interruption of the ascendant water flow by air embolism or xylem occlusion (Zimmermann, 1979; Tyree and Sperry, 1989). Vascular pathogens, which infect the xylem network (Yadeta and Thomma, 2013), are also important drivers of pathogen-mediated plant dieback (Goberville *et al*., 2016; Pandey *et al*., 2018; Fallon *et al*., 2020).

Vascular pathogens induce wood necrosis, leaf symptoms, and crown defoliation (Beckmann and Roberts, 1995; Pearce, 1996). Their biology and toxic metabolite production has been well studied, in particular using controlled phytotoxicity assays (Andolfi *et al*., 2011; Akpaninyang and Opara, 2017). However, the possible role of hydraulic failure during pathogen-mediated plant dieback has been poorly investigated, and the underlying physiological mechanisms inducing leaf symptoms are not clear yet (Fradin and Thomma, 2006; McDowell *et al*., 2008). Moreover, the long-term impact (over seasons) and relationships between pathogens, leaf symptom presence, and the hydraulic functioning of the plant are still unknown. During vascular pathogenesis, both air (Pérez-Donoso *et al*., 2016) and nongaseous (Sun *et al*., 2013; Czemmel *et al*., 2015, Pouzoulet *et al*., 2019) embolism have been observed. For example, air embolism is thought to accelerate pathogen progression during Pierce’s disease (Pérez-Donoso *et al*., 2016), and nongaseous embolism is associated with occlusion of the xylem conduits by the plant that could slow the disease process while interfering with xylem water transport (Sun *et al*., 2013; Pouzoulet *et al*., 2019).

Xylem occlusion, usually through the production of tyloses and gels, is one of the first plant defense mechanisms against vascular pathogens (Pearce, 1996). Xylem parenchyma cells secrete gels and expand into the vessel lumen, forming tyloses, physically blocking pathogen progression (Zimmermann, 1979). Xylem anatomy plays an important role, both for vascular pathogen development (Martin *et al*., 2009; Martín *et al*., 2013; Venturas *et al*., 2014; Pouzoulet *et al*., 2017; 2020) and for tylose formation (Bonsen and Kucera, 1990; De Micco *et al*., 2016; Pouzoulet *et al*., 2019). If effective, this occlusion mechanism allows the plant to compartmentalize the infected zone and to generate new tissue around it (CODIT model, Pearce, 1996). Because tyloses can potentially interfere with the hydraulic functioning of the plant, they could exacerbate disease symptoms (Talboys, 1972). Tyloses are usually observed in close proximity to pathogens, as shown in artificial inoculation studies (Czemmel *et al*., 2015; Rioux *et al*., 2018, among others). However, pathogens frequently proliferate in perennial organs without physically reaching the leaves, thus leaf symptoms are often induced at a distance (Beckmann and Roberts, 1995). A recent study shows that tyloses can be present in symptomatic leaves at a distance from the pathogen niches resulting in decreased leaf hydraulic conductivity (Bortolami *et al*., 2019).

Over the last decades, grapevine (*Vitis vinifera* L.) mortality and yield loss have been reported in European, American, and South African vineyards due to esca trunk disease (Cloete *et al*., 2015; Guerin-Dubrana *et al*., 2019). Esca, a vascular disease caused by the infection of multiple fungal pathogens, affects mostly mature grapevines (more than seven-years-old), and symptoms include trunk necrosis and leaf symptoms, consisting of “tiger-stripe” necrosis and leaf wilting (Lecomte *et al*., 2012; Claverie *et al*., 2020), which are not regularly expressed season-to-season even within individual vines (Guerin-Dubrana *et al*., 2013; Li *et al*., 2017). While the pathogens responsible for esca-induced trunk necrosis have been identified (Morales-Cruz *et al*., 2018; Brown *et al*., 2020), the underlying mechanisms of leaf and fruit symptoms, and plant death are still poorly understood. Bortolami *et al*. (2019) demonstrated that the two vascular pathogens related to esca (*Phaeomoniella chlamydospora* and *Phaeoacremonium minimum)* were never detected in leaves or in stems of the current year, but always in the trunk (independently from leaf symptom presence). They further showed that esca symptomatic leaves presented significant losses in hydraulic conductivity due to the occlusion of the xylem conduits by tyloses. Together, these results reveal that esca impacts leaf hydraulic functioning, but whether or not there is a corresponding failure in perennial organs, and the exact timing of this phenomenon, are still unknown. As stems and branches are the direct connections between the pathogen niche in the trunk and the observed symptoms in the leaves, the study of stem xylem integrity is crucial in the understanding of esca impact on grapevine physiology in the current year and across seasons.

In this study, we investigated stem xylem integrity in grapevine during esca leaf symptom formation asking the following questions: (i) Can esca lead to hydraulic failure in perennial organs? (ii) Does stem hydraulic failure occur prior to or after leaf symptom expression, and does it depend on xylem anatomy? (iii) Do long-term symptomatic plants present different xylem anatomy and levels of hydraulic failure from long-term asymptomatic plants? To answer these questions, we transplanted 28-years-old grapevines (*Vitis vinifera* L. cv Sauvignon blanc) from the field into pots to transport, manipulate, and study naturally esca-infected vines. We coupled *in vivo* visualizations of stem xylem functionality (using synchrotron-based X-ray microcomputed tomography) with stem specific hydraulic conductivity measurements (*k_s_*), theoretical hydraulic conductivity estimates (*k_th_*), optical observations of vessel occlusions, and pathogen detection during symptom appearance, while comparing plants with different symptom history record.

## MATERIALS AND METHODS

### Plant material

*Vitis vinifera* cv. Sauvignon blanc grafted onto 101-14 MGt were uprooted in winter 2017, 2018, and 2019 from a vineyard planted in 1992 located at INRAE Bordeaux-Nouvelle Aquitaine (44°47’24.8”N, 0°34’35.1”W) and transferred into pots. Following plant excavation, the root system (around 0.125 m3) was immersed under water overnight, and powered with indole-3-butyric acid. The plants were potted in 20l pots in fine clay medium (Klasmann Deilmann substrate 4:264) and placed on heating plates at 30 °C for two months. Plants were then moved to a greenhouse, under natural light conditions, and watered with nutritive solutions (0.1 mM NH4H2PO4, 0.187 mM NH4NO3, 0.255 mM KNO3, 0.025 mM MgSO4, 0.002 mM Fe, and oligo-elements [B, Zn, Mn, Cu, and Mo]) until the end of experiment. Since the plantation these plants have been trained with a double Guyot system. This training system requires a permanent main trunk and one cane on each side of the trunk which is left every year to carry the buds that will produce the stems of the year. During the growing season, the stems of the current year were trimmed at 1.5-2m, the secondary stems and inflorescences were removed just after bud-break. Each of these plants has been surveyed each year in the field since 2012 for esca leaf symptom expression following Lecomte *et al*. (2012), and has been classified yearly as leaf-symptomatic or asymptomatic. Plants were then classified by their long-term symptomatology record: plants asymptomatic from 2012 to 2018 (pA, previously asymptomatic), and plants that have expressed symptoms at least once between 2012 and 2018 (pS, previously symptomatic).

### Esca symptom notation

The evolution of esca leaf symptoms was surveyed twice a week from June to October 2019 on every plant (n=84, Fig. 1). As presented in Fig. 1A, esca symptoms were scored at the stem and whole plant scales. The stems of the current year collected for analyses (both hydraulic measurements or microCT observations) could be noted as: asymptomatic (green leaves and apparently healthy), pre-symptomatic (leaves presenting yellowing or small yellow spots between the veins), tiger-stripe (typical pattern of esca leaf symptoms), or apoplectic (leaves passing from green to wilted in a couple of days). Along the experimentation, entire plants could be noted as asymptomatic (control) or symptomatic (when at least 25% of the canopy was presenting tiger-stripe leaf symptoms). At the end of the experiment (week 40, October 2019) each plant was classified as symptomatic or asymptomatic (control). We were then able to group each stem measured into six different groups (Fig. 1A): one group of stems from control plants (asymptomatic from June to October) and five groups of stems from symptomatic plants: two before symptom appearance (asymptomatic and pre-symptomatic stems); and three after symptom appearance (asymptomatic, tiger-stripe, and apoplectic stems). To clearly differentiate asymptomatic stems collected from symptomatic plants and asymptomatic stems collected from asymptomatic plants, we considered plants (and their stems) that didn’t show leaf symptoms during the experiment as control plants (or stems). We investigated whether symptom expression (final symptom notation in October 2019, see Fig. 1) differed between plants with contrasted long-term symptom history (previously asymptomatic vs previously symptomatic, Table 1) using a Chi-square test of independence.

**Fig. 1.**
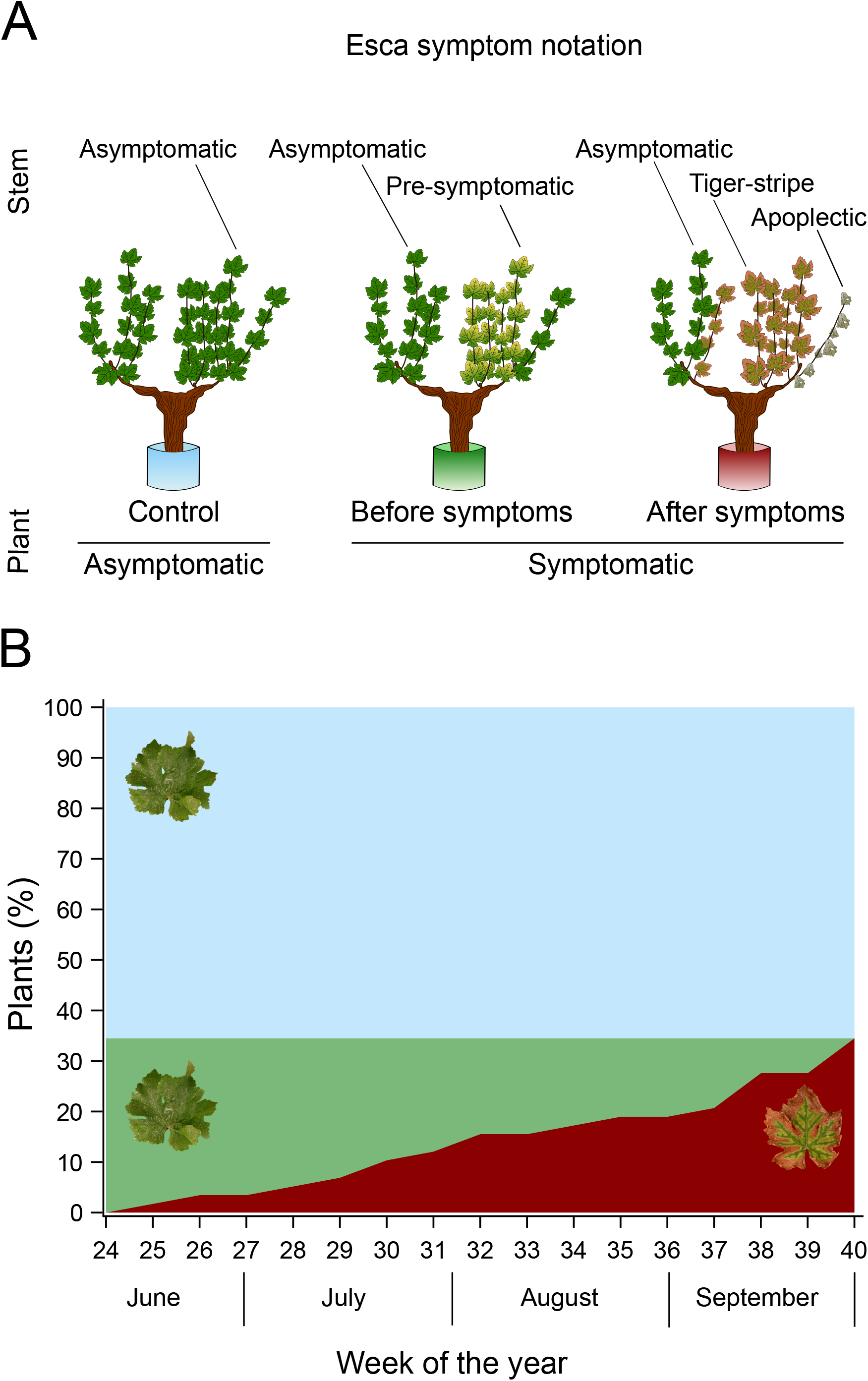
Representation of esca symptom notation during the experimental season. (**A**) Single stems could be noted as esca asymptomatic, pre-symptomatic, tiger-stripe, or apoplectic. Whole plants have been noted as control (asymptomatic from June to October) or symptomatic (with tiger-stripe symptoms at the end of the season). (**B**) Proportion of plants in each symptom category over the experimental season (n=58). The blue area corresponds to control plants, green area to esca symptomatic plants before symptom appearance, and red area to esca symptomatic plants.

**Table 1.** Esca leaf symptom observations over the experimental season on *Vitis vinifera* cv Sauvignon blanc.

| Symptom notation<br>before 2019 | All plants | Previously<br>asymptomatic (pA) | Previously<br>symptomatic (pS) |
| --- | --- | --- | --- |
| **Symptom notation in 2019** |  |  |  |
| Esca-symptomatic | 35 % (20/58) | 30 % (6/20) | 37 % (14/38) |
| Control-asymptomatic | 65 % (38/58) | 70 % (14/20) | 63 % (24/38) |

Plants are grouped by their symptom history: previously asymptomatic (pA, plants that have never expressed leaf symptoms between 2012 and 2018) and previously symptomatic (pS, plants that have expressed leaf symptoms at least once since 2012). Ratios present the number of plants in each symptom category (esca-symptomatic or control-asymptomatic) over the total number of plants of the category.

### X-ray microCT observation

Synchrotron-based microCT was used to visualize the content of vessels and their functionality in esca tiger-stripe and control stems. Three symptomatic plants (presenting tiger-stripe symptoms for 8, 7, and 3 weeks), and one asymptomatic-control plant were brought to the PSICHE beamline (King *et al*., 2016) at SOLEIL synchrotron facility in September 2019. Stems of the current year (ca. 2 m long) were cut under water and transferred into a solution containing 75mM of contrasting agent iohexol. The iohexol solution absorbs X-rays very strongly and appears bright white in X-ray scans above the iodine K-edge at 33.2 keV, and, once it has been taken up by the transpiration stream, the effective functionality of each vessel can be confirmed (Pratt and Jacobsen, 2018; Bortolami *et al*., 2019). These stems were moved and left outdoor to transpire the solution for at least half a day. The stems were then transferred to the beamline stage and scanned twice in less than 5 minutes using two different energies of a high-flux (3 x 10^11^ photons mm^-2^) monochromatic X-ray beam: 33.1 keV and 33.3 keV. The projections were recorded with a sCMOS camera equipped with a 250-mm-thick LuAG scintillator (Orca Flash, Hamamatsu, Japan). The complete tomographic scan included 1500 projections, and each projection lasted 50 ms. Tomographic reconstructions were performed using PyHST2 software (Mirone *et al*., 2014) using the Paganin method (Paganin *et al*., 2002), resulting in 32-bit volume reconstructions of 2048 x 2048 x 1024 voxels. The final spatial resolution was 2.8769 μm^3^ voxel^-1^.

### Image analysis of microCT scans

The contrast agent iohexol allowed us to distinguish in intact scans the effective functionality of each vessel. In the absence of iohexol, X-ray microCT scans are used to distinguish air-filled vessels (appearing black, corresponding to native PLC) from sap-filled vessels (appearing grey). The addition of iohexol in the xylem sap allows to distinguish the functional vessels (they appear bright white when they transport the sap), from the non-functional ones (i.e. occluded vessels remaining grey, corresponding to occlusion PLC). We could also observe partially occluded vessels (i.e. vessels with simultaneous presence of air and occlusions, or sap and occlusions). This specific case was observed by checking the presence of any occlusion in at least 200 slices in each volume. Partially occluded vessels were considered as occluded, some examples are presented in Fig. S1. The equivalent-circle diameter of air-filled, occluded, and functional (iohexol-filled) vessels was measured on the cross sections from the central slice of the microCT scanned volume using ImageJ software (Schneider *et al*., 2012). In the high energy scans recorded at 33.3 keV X-ray beam, iohexol appears bright white but its contrast can sometimes impede the clear limit of the vessel lumen. Therefore, all vessel diameters were recorded on the scan recorded at low energy (33.1 keV X-ray beam), then the distinction of occluded from iohexol-filled vessels was done on the high energy scan (as done by Bortolami et al. 2019). The theoretical hydraulic conductivity of each vessel (*k_vessel_*) [kg m MPa^-1^ s^-1^] was calculated using the Hagen-Poiseuille equation:

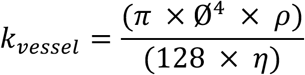

Where: *Ø* is the equivalent circle diameter [m], *ρ* the density of water [998.2 kg m-3 at 20°C], and *η* the viscosity of water [1.002 x 10-9 MPa s at 20°C]. The percentage loss of hydraulic conductivity given by native air embolism (native PLC) was calculated by the ratio between the hydraulic conductivity of air-filled vessels and the whole-stem hydraulic conductivity:

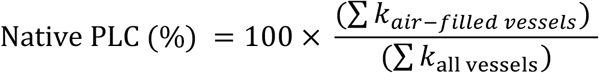

The percentage loss of hydraulic conductivity given by occlusions (occlusion PLC) was calculated by the ratio between occluded (plus partially occluded) vessels and the whole-stem hydraulic conductivity:

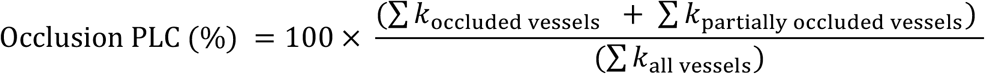

The total percentage loss of hydraulic conductivity (total PLC) was obtained by summing native PLC with occlusion PLC in each sample. As the first ring of xylem vessels (i.e. protoxylem) was always non-functional (>90% PLC), both in control and tiger-stripe stems, it was removed from the analysis.

We investigated whether native PLC, and occlusion PLC differed between control and esca tiger-stripe plants, using two independent generalized mixed linear models where plants were treated as a random effect. Proportional data (ranging from 0 to 1, dividing all PLC values by 100) was analyzed to fit a logit link function and binomial distribution as appropriate.

### Monitoring stem hydraulic properties over time

Xylem integrity was monitored over time by measuring hydraulic properties in stems produced on the year of the experiment and collected on control and symptomatic plants along the season and during esca development. Specific hydraulic conductivity (*k_s_*) was measured on internodes sampled in the center of the collected stem by the gravity method (Sperry *et al*., 1988), and compared to its theoretical analog (*k_th_*) calculated from xylem anatomical observations on the same internode or on the one below (see the method described below). When there are observed differences in *k_s_* among stems, comparisons with theoretical maximums (*k_th_*) can show if lower *k_s_* values result from anatomical differences (i.e. different vessel size distributions) or by hydraulic failure (in the case of similar vessel size and density). If *k_s_* varies in unity with *k_th_*, differences in *k_s_* might result from anatomical differences (e.g. smaller *k_s_* are related to smaller vessels), otherwise *k_s_* variations are the consequence of hydraulic failure. Each method to measure *k_s_*, *k_th_*, and to observe tyloses is described below.

Sampling started on June 19^th^ and finished on September 13^th^ 2019 for a total of 10 sampling dates, 39 stems of the current year from 23 control-asymptomatic plants, and 49 stems of the current year from 17 symptomatic plants. We randomly sampled control plants and esca symptomatic plants all along the season through the evolution of esca symptoms, obtaining measurements from 14 weeks before until 10 weeks after symptom appearance. To explore the contribution of the experimental design to data analysis, we tested the effect of the year of uprooting (2018 and 2019), the position of the analyzed internode, and the week of the measurement (i.e. evolution during the season) on *k_s_* and *k_th_* in control plants using separate generalized linear mixed model with normal distributions and the plant treated as a random variable (Table S1). A significant impact of the year of uproot was found for *k_s_* and *k_th_* values in control plants (Table S1). This could have resulted from the more favorable conditions (i.e. climatic stability and nutrient availability) for the greenhouse grown vines (note that plants uprooted in 2017 were only esca symptomatic and were not included in this analysis). However, once *k_s_* and *k_th_* are plotted together (Fig. S2), all the values lie on the same regression line without generating outlier values (smaller *k_s_* values correspond to smaller *k_th_* values independently of the uprooting year).

### Stem specific hydraulic conductivity (k_s_)

*k_s_* measurements were performed on one internode per stem, located in the center of the collected >1.5m long stem, following Torres-Ruiz *et al*. (2012) gravity method. In the early morning, each stem was cut at the base under water to avoid air entrance in the stem, maintained under water and brought to the laboratory. Hydraulic conductivity measurements were always done before noon, in order to minimize the delay (never more than four hours) from the cut to the measure. In the laboratory, a representative internode between the 4th to the 10th internode from the base (i.e. in the center of the stem) was cut underwater with a clean razor blade, the ends wrapped in tape, and the internode was connected to a pipe system. A flow of 20 mM KCl solution passed through the sample from a reservoir to a precision electronic balance (AS220.R2, RADWAG, Radom, PL) recording the weight every 5 seconds using the WinWedge v3 5.0 software (TAL Technologies, Philadelphia, PA, USA). The solution was passed through the stem at four increasing pressures (ranging from 0.001 to 0.005 MPa), controlled by raising the source height. The average flow for each pressure step was determined after stabilization at a steady-state as the average of 10-15 measures. Hydraulic conductance, *k* [kg s^-1^ MPa^-1^] was obtained by the slope generated by the flow and the corresponding pressure. The linear relationship between flow and pressure obtained were always characterized by R^2^>0.97. Stem specific hydraulic conductivity, *k_s_* [kg s^-1^ MPa^-1^ m^-1^], was calculated as follows:

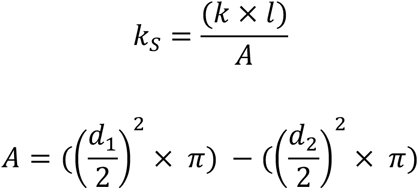

Where: *k* is the hydraulic conductance, *l* is the length of the sample, *A* is the xylem area, *d_1_* is the external diameter of the debarked stem, and *d_2_* is the diameter of the central pith.

### Stem theoretical hydraulic conductivity (*k_th_*), vessel anatomy, and tylose observation

Just before hydraulic conductivity (*k_s_*) measurements, the lower internode was stored at 4 °C in 80% ethanol for analysis of xylem anatomy. When possible, the same internode of *k_s_* measurements was used for anatomical analysis and *k_th_* estimations, otherwise the stored internode was used for the following protocol. 50 μm thick slices were obtained using a GSL-1 microtome (Gärtner *et al*., 2014). Slices were stained using a 0.5% safranin solution during 5 minutes, and then washed three to four times in ethanol (100%). They were quickly soaked in xylene and mounted on microscope slides with Permount Mounting Medium (Electron Microscopy Science, Hatfield, PA, USA). Images were captured with a stereo microscope SMZ1270 (Nikon, France) mounted with a DS-Fi3 camera (Nikon, France). The theoretical conductivity of each vessel (*k_vessel_*) [kg m Mpa^-1^ s^-1^] was calculated using the Hagen-Poiseuille equation as described above.

Where *Ø* is the equivalent circle diameter [m] (measured with ImageJ software), *ρ* the density of water [998.2 kg m^-3^ at 20 °C], and *η* the viscosity of water [1.002 x 10^-9^ MPa s at 20 °C]. *k_th_* of the stem [kg s^-1^ m^-1^ Mpa^-1^] was then calculated by summing every *k_vessel_* in the xylem area (*A*) [m^2^]:

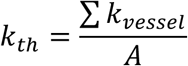

In the entire cross section of each sample, the physical presence (or absence) of tyloses in vessel lumina was visually assessed.

Regarding the statistical analysis, stems were grouped in six different categories following their esca symptomatology (as presented in Fig. 1A). We investigated whether *k_s_*, *k_th_*, and total vessel density differed among these different categories, and how *k_s_*, *k_th_*, and total vessel density differed between stems with and without tyloses (independently from leaf symptom presence), using independent mixed linear general models. The symptom / tylose category and the year of uprooting (since it had a significant impact on *k_s_* and *k_th_* in control plants, Table S1) were entered as fixed effects, with the plant treated as a random effect since different stems were sometimes analyzed from the same plant (88 analyzed stems on 40 different plants). Total density and densities for each vessel diameter class were log-transformed prior to analysis to fit normality requirements. For the classes with no vessels (e.g. samples without vessel diameters above 160 μm), a minimal density of 0.0001 was assigned prior to log transformation. We investigated whether the frequency of symptomatic stems presenting tyloses changed with the symptom age (i.e. weeks between first symptom detection and *k_s_* measurements on the same plant) with a Chi-square test. The relationships between stem *k_s_* and *k_th_* were tested using linear regression models. Finally, we investigated whether *k_s_* and *k_th_* in control stems differed between plants with different symptom history records using independent mixed linear general models with the plant treated as a random effect.

### Fungal detection

Detection and quantification of *Phaeomoniella chlamydospora* and *Phaeoacremonium minimum* were performed using qPCR in a subsample of stems of the current year (n = 28) and perennial trunks (n = 20 plants) from the same symptomatic and control plants used for hydraulic and anatomical measurements. All along the season, basal internodes, from the same stems sampled for *k_s_* and *k_th_* measurements, were directly placed in liquid nitrogen and stored at −80 °C. At the end of the experiment, a subset of plants was cut at the base for trunk sampling. A 2 cm high section was cut with a sterilized hand saw. The bark was removed and the different tissues of each section (necrotic and apparently healthy wood) were separately collected using ethyl alcohol-sterilized shears in a sterile environment, and immediately placed in liquid nitrogen. All samples were ground in liquid nitrogen using a tissue lyser (Tissuelyser II, Qiagen, Germantown, MD, USA). DNA was extracted from 60mg of ground tissue using the Invisorb Spin Plant Mini Kit (Invitek GmbH, Berlin, Germany)according to the manufacturer’s instructions. Detection and quantification of *P. chlamydospora* and *P. minimum* (previously named *P. aleophilum*) DNA by qPCR (SYBR Green assays) was conducted using the primer sets PchQF (5’-CTCTGGTGTGTAAGTTCAATCGACTC-3’)/PchQR (5’-CCATTGTAGCTGTTCCAGATCAG-3’) and PalQF(5’-CCGGTGGGGTTTTTACGTCTACAG-3’)/ PalQR(5’-CGTCATCCAAGATGCCGAATAAAG-3’) (Pouzoulet *et al*. 2013). The qPCR reactions proceeded in a final volume of 25 μl, and the reaction mixtures containing 2 μL of DNA template, 12.5 μl of 2X SYBRGreen Quantitect Master Mix (Qiagen, Venlo, Netherlands), and each primer at a final concentration of 0.4 μM. Experiments were conducted with a Mx3005P Real-Time PCR cycler using MxPro qPCR software (Agilent Technologies). The cycling program, as described in Pouzoulet *et al*. (2017), consisted of an initial denaturation step at 95°C for 15 min, and 40 cycles of 15 s at 95°C (for denaturation) followed by 45 s at 62°C (for both annealing and extension). A melting analysis of 40 min from 60 to 95° was performed to verify reaction’s specificity and the absence of byproducts. Preparation and use of standard solutions for the absolute quantification of fungal DNA was realized following Pouzoulet *et al*. (2013) using ten-fold dilutions of fungal DNA extracts obtained from axenic cultures. Reaction efficiencies ranging from 90% and 95% with an R^2^ > 0.99 (n=15) were obtained for both PchQF/R and PalQF/R primer sets. The average amount of DNA was determined based on three technical replicates (standards and plates) with a detection threshold superior to 95% (i.e. at least three positive amplification out of three replicates) or otherwise discarded (i.e. pathogen DNA was considered absent). Pathogen DNA quantity (average value of three technical replicates, fg/μl) was normalized by the amount of total DNA (ng/μl), measured using a Qubit fluorometer. The results from each trunk sample (i.e. necrotic or apparently healthy wood) were averaged together in order to obtain one quantification per plant. We investigated whether the amount of fungal DNA (both for *P. chlamydospora* and for *P. minimum*) in trunks differed between symptomatic and control plants, and between control plants with different symptom history records, using generalized linear mixed model with a poisson distribution and a log likelihood function.

### Statistical analysis

All data management and statistical tests were done in SAS software (SAS 9.4; SAS Institute). We used PROC GLIMMIX for generalized linear mixed models, PROC GLM for generalized linear models, PROC REG for regression analyses and PROC FREQ for frequency analyses (Chi-square test of independence). The normality of the response variables was tested using a Kolmogorov-Smirnov test (PROC UNIVARIATE) prior to analyses. Data were log-transformed (total density) or appropriate distributions (binomial, poisson) were fitted when appropriate.

## RESULTS

### Esca leaf symptom expression within and across seasons

Esca leaf symptoms were recorded in 20 out of the 58 plants followed in this study (35%, Fig. 1, Table 1). The number of symptomatic plants increased gradually with time, from the first symptom appearance in early June to the last in late September (Fig. 1). There was no effect of the plant history (previously asymptomatic pA, or previously symptomatic pS) on 2019 symptom expression (n=58, *X*^2^=0.27, P=0.60). On 20 pA plants, six (30%) expressed leaf symptoms in 2019 (Table 1). On 38 pS plants, fourteen (37%) showed symptoms in 2019 (Table 1). However, pS plants expressed symptoms from June to the end of September, while pA plants showed leaf symptoms only in September.

### *In vivo* observations of esca symptomatic stems

Xylem vessels of control and tiger-stripe stems were observed using three dimensional X-ray microCT scans in iohexol-fed samples (Fig. 2, 3, Table S2). As shown in Fig. 2, functional and non-functional vessels can be discriminated through the use of iohexol (functional vessels appear bright white, non functional vessels appear either black if air-filled or grey if occluded).

**Fig. 2.**
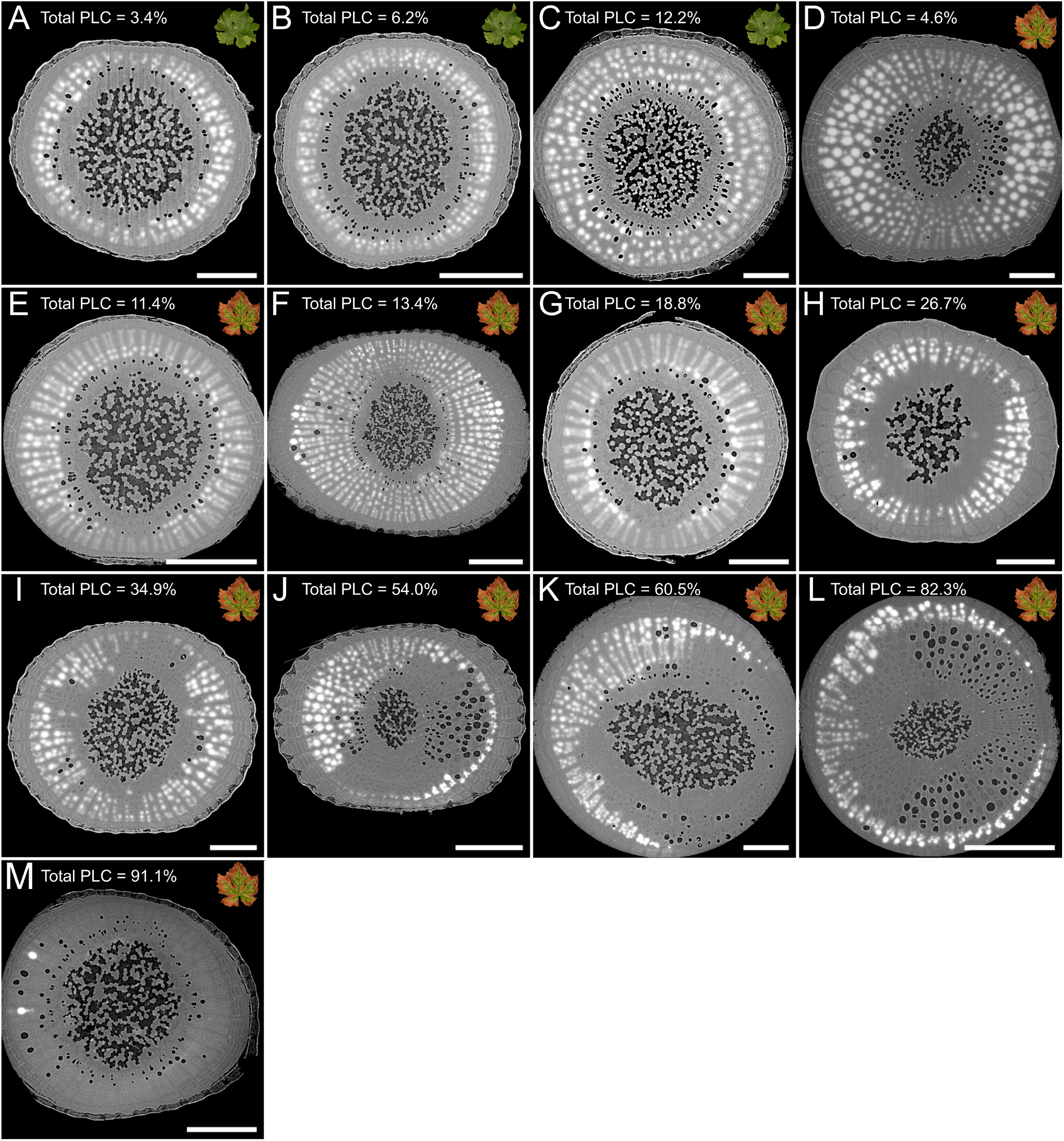
Two-dimensional reconstruction of cross-sections from X-ray microCT volumes of grapevine stems. Each panel represents a cross section of different stems for control (**A-C**) and esca symptomatic (**D-M**) plants. Iohexol appears white bright in functional vessels; air-filled vessels (i.e. native PLC) appear black; occluded vessels (i.e. occlusion PLC) appear grey. Total PLC (i.e. native PLC plus occlusion PLC) values are given for the presented samples. Scale bars = 1000μm.

**Fig. 3.**
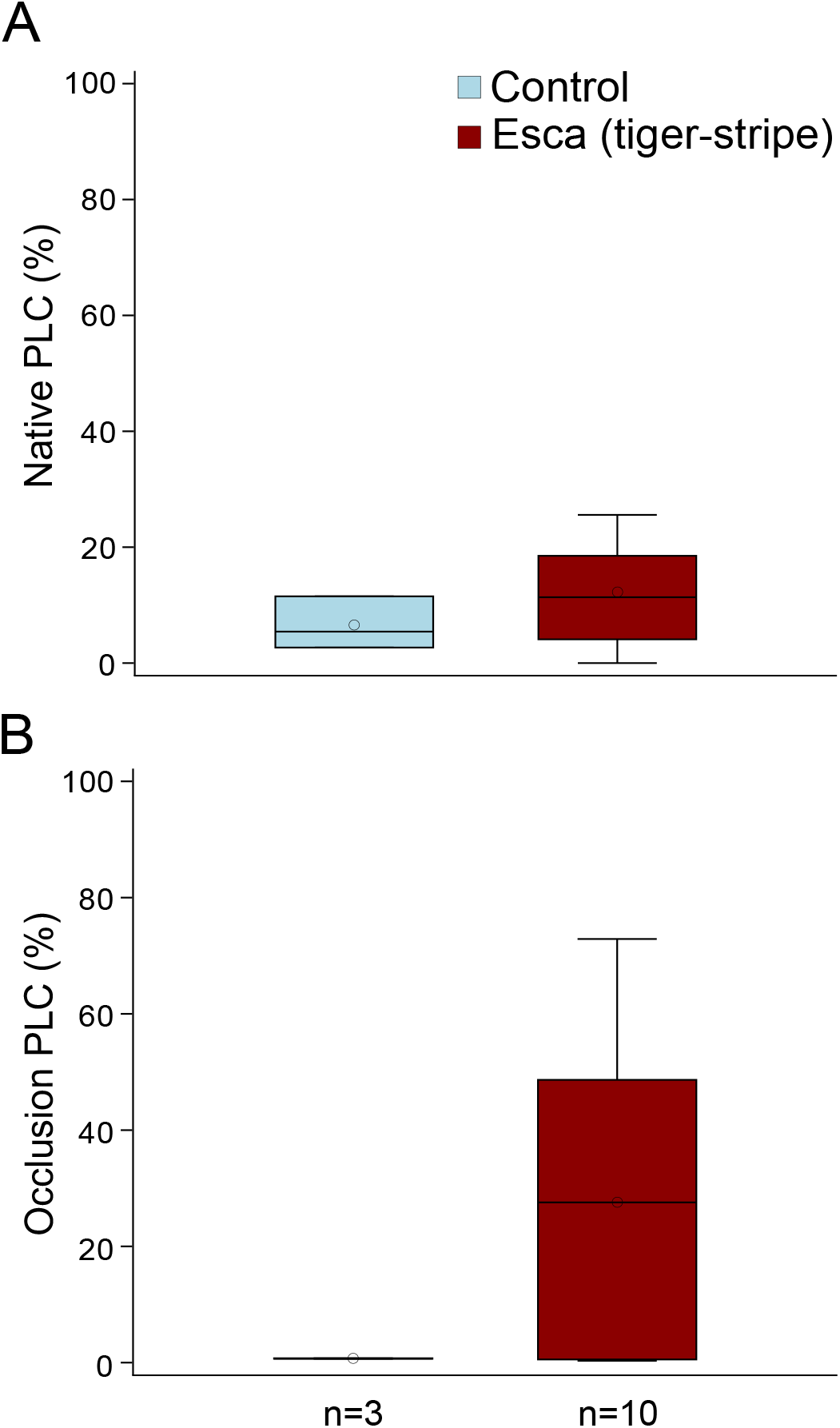
(**A**) Mean values of native PLC in control (blue) and esca tiger-stripe (red) stems of grapevine plants using X-ray microCT imaging. Differences were not significant (n=13, F_1,9_=0.07, P=0.79). (**B**) Mean values of occlusion PLC in control (blue) and esca tiger-stripe (red) stems of grapevine plants using X-ray microCT imaging. Differences were not significant (n=13, F_1,9_=0.33, P=0.58). Boxes and bars show the median, quartiles and extreme values, circles show mean values. N represents the sample size (number of analyzed stems) for each group.

We observed almost totally functional stems in all asymptomatic stems (<20% total PLC, Fig. 2A-C), and 40% of tiger-stripe stems (e.g. Fig. 2D-G). Higher levels of PLC (>20% total PLC, Fig H-M) were observed in the remaining tiger-stripe stems, with 40% of tiger-stripe stems exhibiting over 50% total PLC (Fig. 2J-M). When the two components of PLC were disentangled, we observed that the level of native PLC remained low both in control (6.5 ± 2.6%) and in tiger-stripe (12.2 ± 2.9%) stems (Fig. 3A). Occlusion PLC values were virtually zero in control stems (0.7 ± 0.02%) while in tiger-stripe stems the mean occlusion PLC values was 27.5 ± 8.2% (Fig. 3B). Nevertheless, the variability of occlusion PLC across tiger-stripe stems was very high, the values ranging from 0.3% to 72.9% (Fig. 2D-M, and 3B), and occlusion PLC was not correlated to symptom age (n=10, F_2,7_=0.19, P=0.83). Consequently, no statistical differences in native or occlusion PLC were found between control and tiger-stripe stems (Fig. 3). When higher occlusion PLC was measured (Fig. 2H-M), occluded vessels could be organized either on one side of the stem (Fig 2J-L) or randomly distributed across the section (Fig 2H, 2I, 2M). In 90% of symptomatic stems, we observed that the most external vessels were functional. Occlusions were present equally in all vessel diameter classes (Fig. S3).

### Tylose development, stem specific (*k_s_*) and theoretical (*k_th_*) hydraulic conductivity during esca leaf symptom formation

Tyloses were identified in the xylem vessels of certain tiger-stripe stems and throughout the temporal development of esca leaf symptoms, from the appearance of symptoms to 11 weeks after. All apoplectic stems and 62.5% (15 of 24 analyzed stems) of esca tiger-stripe stems presented tyloses, while all other stems (control, asymptomatic or pre-symptomatic) did not contain these occlusions, even until one week before symptom development. In esca tiger-stripe stems, tyloses were not related to specific plants, or to symptom age (i.e. on the same plant at the same moment, different symptomatic stems could present tyloses, or not, n=24, *X*^2^=7.47, P=0.38).

Overall, no significant impact of esca symptoms was observed on *k_s_* (Fig. 4A), even if tiger-stripe stems were divided between those with and without tyloses. Control stems presented a mean (± SE) *k_s_* of 24.97 ± 1.72 kg s^-1^ MPa^-1^ m^-1^; all the stems without tyloses measured on symptomatic plants showed the same range of values as control stems (Fig 4A, Table 2): 26.04 ± 4.71 for asymptomatic before symptoms appearance, 30.32 ± 4.26 for pre-symptomatic stems, 19.80 ± 5.18 for asymptomatic stems after symptom appearance on the plant, and 21.29 ± 5.40 for tiger-stripe stems without tyloses. Stems with tyloses (tiger-stripe and apoplectic stems) presented the lowest average *k_s_* values (11.27 ± 2.86 and 2.47 ± 1.45 kg s^-1^ MPa^-1^ m^-1^ for tiger-stripe and apoplectic, respectively). Regarding *k_th_*, no significant impact of esca symptoms was found (Fig. 4C, Table 2), all the values were in the same range, with average values ranging from 70.44 (for tiger-stripe stems with tyloses) to 87.88 (for pre-symptomatic stems) kg s^-1^ MPa^-1^ m^-1^.

**Fig. 4.**
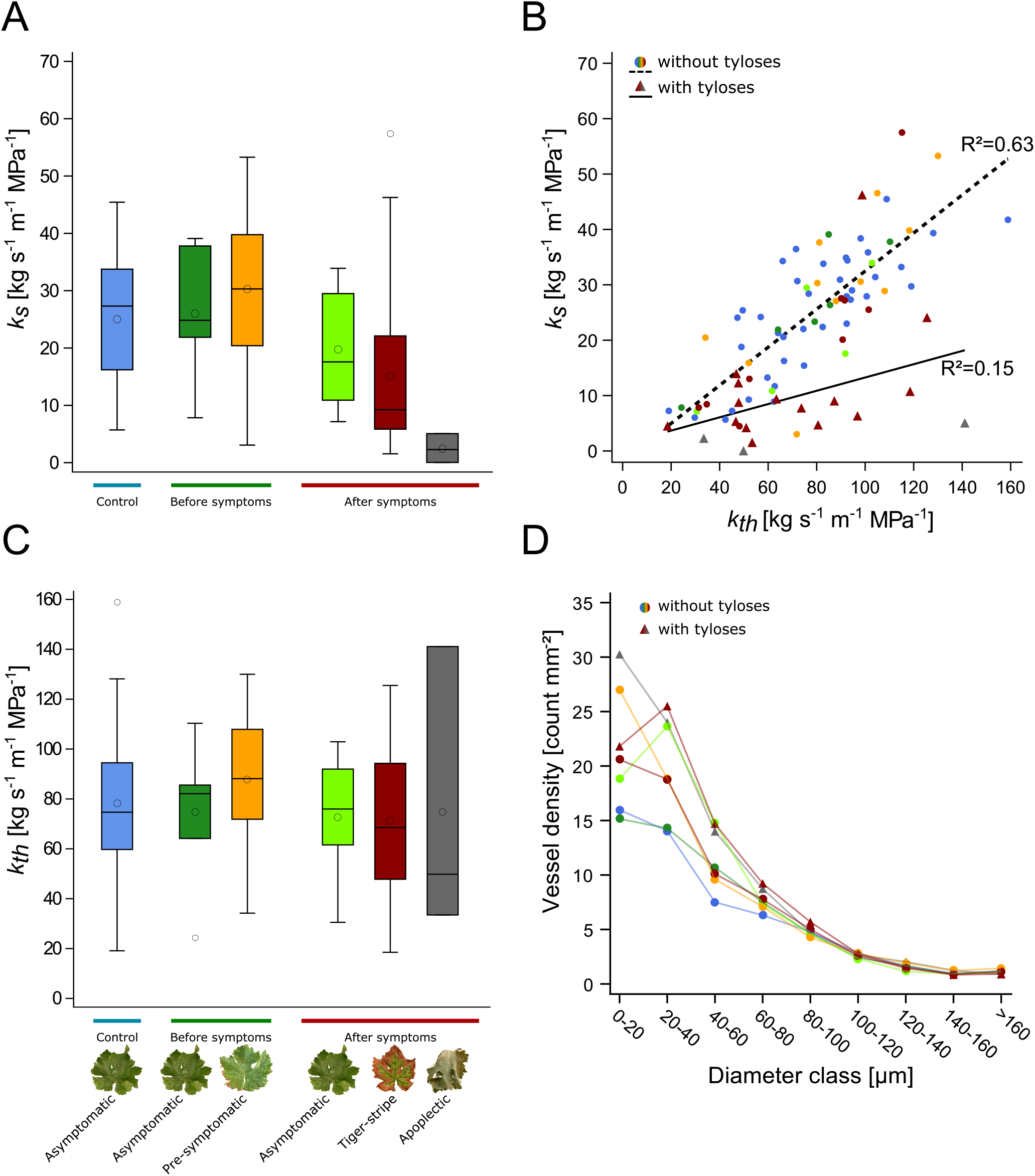
Relationships between specific stem hydraulic conductivity (*k_s_*) theoretical stem hydraulic conductivity (*k_th_*), and vessel density in control and esca symptomatic grapevine plants. (**A**) *k_s_* values for control (blue); asymptomatic (dark green) and pre-symptomatic (yellow) stems in plants before symptom appearance; asymptomatic (light green), tiger-stripe (red), and apoplectic (grey) stems in plants after symptom appearance, differences were not significant (n=88, F_5,45_=1.30 P=0.28). Boxes and bars show the median, quartiles and extreme values, circles within boxes correspond to means, and circles outside boxes to outlier values. (**B**) Relationships between *k_s_* and *k_th_*. Symbols represent the absence (circles) or presence (triangles) of tyloses in xylem vessels. Colors represent esca symptomatology (as in panel A). The dashed line represents the regression for stems in which no tyloses were observed in xylem vessels, and the solid line represents the regression for samples with tyloses. R^2^ for the regression lines are indicated (see Table 2 and Fig. S4 for detailed analyses). (**C**) *k_th_* values for the different stem categories as presented in panel a. Differences were not significant (n=88, F_5,45_=0.58, P=0.71). (**D**) Relationships between mean values of xylem vessel density and their diameters. Differences in total vessel density and in vessel size distributions were not significant (n=88, F_6,45_=0.77, P=0.60; n=792 (88 samples for 9 vessel classes), F_48,693_=1.19, P=0.18). Colors and markers are the same as panel B.

**Table 2.** Values for specific stem hydraulic conductivity (*k_s_*), theoretical stem hydraulic conductivity (*k_th_*) and equations of regression lines between *k_s_* and *k_th_* for control and esca symptomatic stems.

| Tyloses | Esca | $k_s$<br>[kg s<sup>-1</sup> m<sup>-1</sup> MPa<sup>-1</sup>] | $k_{th}$<br>[kg s<sup>-1</sup> m<sup>-1</sup> MPa<sup>-1</sup>] | n<br>(stem - plant) | Regression |
| --- | --- | --- | --- | --- | --- |
| Absence | Control | 24.97 ± 1.72 | 78.36 ± 4.51 | 39 - 23 | $k_s = 0.3 \times k_{th} + 1.6$<br>R<sup>2</sup>=0.61 P<0.0001 |
| | Asymptomatic before symptoms | 26.04 ± 4.71 | 74.75 ± 11.80 | 6 - 6 | $k_s = 0.36 \times k_{th} - 0.94$<br>R<sup>2</sup>=0.82 P=0.013 |
| | Pre-symptomatic | 30.32 ± 4.26 | 87.88 ± 8.54 | 11 - 7 | $k_s = 0.37 \times k_{th} - 1.9$<br>R<sup>2</sup>=0.54 P=0.010 |
| | Asymptomatic after symptoms | 19.80 ± 5.18 | 72.58 ± 12.64 | 5 - 2 | $k_s = 0.33 \times k_{th} - 4$<br>R<sup>2</sup>=0.64 P=0.104 |
| | Esca (tiger-stripe) | 21.29 ± 5.40 | 72.85 ± 10.41 | 9 - 5 | $k_s = 0.45 \times k_{th} - 11.26$<br>R<sup>2</sup>=0.74 P=0.003 |
| Presence | Esca (tiger-stripe) | 11.27 ± 2.86 | 70.44 ± 7.81 | 15 - 5 | $k_s = 0.17 \times k_{th} - 0.84$<br>R<sup>2</sup>=0.22 P=0.077 |
| | Esca (apoplectic) | 2.47 ± 1.45 | 74.80 ± 33.48 | 3 - 2 | $k_s = 0.04 \times k_{th} - 0.2$<br>R<sup>2</sup>=0.68 P=0.385 |
| Absence | All | 25.06 ± 1.46 | 78.42 ± 3.37 | 70 - 37 | $k_s = 0.34 \times k_{th} - 1.90$<br>R<sup>2</sup>=0.63 P<0.0001 |
| Presence | All | 9.81 ± 2.51 | 71.16 ± 8.00 | 18 - 7 | $k_s = 0.12 \times k_{th} - 1.28$<br>R<sup>2</sup>=0.15 P=0.117 |

In order to further investigate the impact of esca on stem hydraulics, we explored the relationship between individual stem *k_s_* and *k_th_* in each symptom category (Fig. 4B, S4, Table 2). Significant relationships were found between *k_s_* and *k_th_* in all groups except in asymptomatic stems after symptom appearance and symptomatic stems with the physical presence of tyloses (Fig. S4, Table 2). The slopes of regression curves between *k_s_* and *k_th_* did not vary among groups in the absence of tyloses (slope values ranged between 0.3 and 0.4, Table 2) while it was close to 0 in the presence of tyloses (0.17 for tiger-stripe and 0.04 for apoplectic stems). When *k_s_* and *k_th_* are compared in the presence or absence of tyloses, we observed that *k_s_* was significantly lower when tyloses were present (9.81 ± 2.51 kg s^-1^ MPa^-1^ m^-1^ in the presence of tyloses vs 25.06 ± 1.46 kg s^-1^ MPa^-1^ m^-1^ in the absence of tyloses, Table 2, n=88, F_1,49_=7.11, P=0.01) while *k_th_* did not significantly differ. Stems without tyloses presented a strong correlation between *k_s_* and *k_th_*, while in the presence of tyloses this relationship was not significant (Table 2, Fig. 4B).

Total vessel density did not significantly differ between stem symptomatology (comparing all the seven categories presented in Table 2), even when vessel density was partitioned by vessel diameter classes (Fig. 4D).

Finally, we tested the impact of disease history (comparing pA and pS plants) on the hydraulic conductivity and xylem anatomy in control plants. There were no differences between longterm symptomatic (pS) and long-term asymptomatic (pA) plants in stem *k_s_*, stem *k_th_*, or total vessel density (Table 3).

**Table 3.** Long-term impact of symptom presence (i.e. comparing plants with different disease history record) in control plants on specific stem hydraulic conductivity (*k_s_*), theoretical stem hydraulic conductivity (*k_th_*), stem total vessel density, and amount of *Phaeomoniella chlamydospora* and *Phaeoacremonium minimum* DNA in trunks of plants without foliar symptoms.

|  | Previously<br>asymptomatic (pA) | Previously<br>asymptomatic (pS) | Type III Tests of<br>Fixed Effects (pA vs pS) |
| --- | --- | --- | --- |
| $k_s$ [kg s<sup>-1</sup> m<sup>-1</sup> MPa<sup>-1</sup>] | 23.76 ± 2.30 | 26.54 ± 2.61 | n=39, F<sub>1,16</sub>=1.19, P=0.29 |
| $k_{th}$ [kg s<sup>-1</sup> m<sup>-1</sup> MPa<sup>-1</sup>] | 72.22 ± 4.85 | 86.30 ± 7.98 | n=39, F<sub>1,16</sub>=3.01, P=0.10 |
| total vessel density<br>[count mm<sup>-2</sup>] | 57.28 ± 4.03 | 52.61 ± 3.25 | n=39, F<sub>1,16</sub>=0.72, P=0.41 |
| *P. chlamydospora*<br>[pg ng<sup>-1</sup>] | 6.14 ± 1.90 | 10.15 ± 3.41 | **n=13, F<sub>1,11</sub>=5900.06,**<br>**P<0.0001** |
| *P. minimum* [pg ng<sup>-1</sup>] | 9.27 ± 6.97 | 26.40 ± 13.83 | **n=13, F<sub>1,11</sub>=51014, P<0.0001** |

Values represent means ± SE. Pathogen quantification was estimated as: pg fungal DNA ng<sup>-1</sup> total DNA. Statistical tests used are individual generalized linear mixed models to compare pA vs pS plants (fixed effect) with the individual plants entered as a random effect in the models and the year of uprooting as a co-variable (fixed effect). Statistically significant results (P<0.05) are shown in bold.

### Fungal detection

The two vascular pathogens associated with esca (*Phaeomoniella chlamydospora* and *Phaeoacremonium minimum*) were never detected in stems of the current year while they were systematically detected in the perennial trunk of both control and symptomatic plants (Table 4). In trunks, a significantly higher quantity of fungal DNA was detected in tiger-stripe symptomatic plants than in controls (Table 4). We found 2.14- and 1.64-fold more of *P. chlamydospora* and *P. minimum* DNA in symptomatic trunks relative to controls. In control plants, different symptom history records impacted the quantity of fungal DNA detected by qPCR, for *Phaeomoniella chlamydospora*, and for *Phaeoacremonium minimum*. We found 1.65- and 2.84-fold more *P. chlamydospora* and *P. minimum* DNA in previously symptomatic trunks relative to previously asymptomatic trunks (Table 3).

**Table 4.**
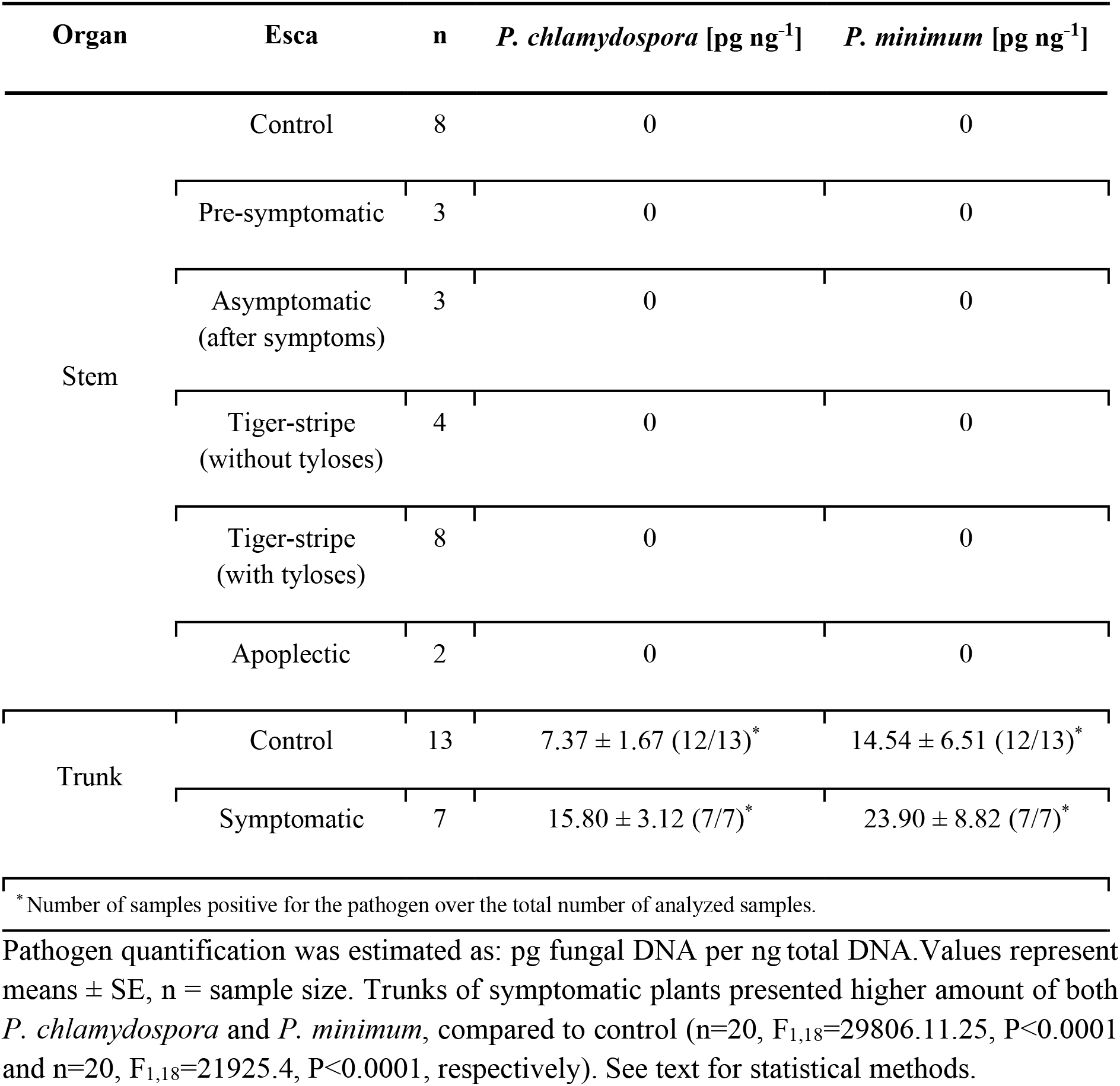
Quantification by qPCR of *Phaeomoniella chlamydospora* and *Phaeoacremonium minimum* DNA in stems and trunks of different esca symptomatology.

| Organ | Esca | n | *P. chlamydospora* [pg ng<sup>-1</sup>] | *P. minimum* [pg ng<sup>-1</sup>] |
| --- | --- | --- | --- | --- |
| Stem | Control | 8 | 0 | 0 |
|  | Pre-symptomatic | 3 | 0 | 0 |
|  | Asymptomatic<br>(after symptoms) | 3 | 0 | 0 |
|  | Tiger-stripe<br>(without tyloses) | 4 | 0 | 0 |
|  | Tiger-stripe<br>(with tyloses) | 8 | 0 | 0 |
|  | Apoplectic | 2 | 0 | 0 |
| Trunk | Control | 13 | 7.37 ± 1.67 (12/13)* | 14.54 ± 6.51 (12/13)* |
|  | Symptomatic | 7 | 15.80 ± 3.12 (7/7)* | 23.90 ± 8.82 (7/7)* |

\*Number of samples positive for the pathogen over the total number of analyzed samples.
Pathogen quantification was estimated as: pg fungal DNA per ng total DNA. Values represent means ± SE, n = sample size. Trunks of symptomatic plants presented higher amount of both *P. chlamydospora* and *P. minimum*, compared to control (n=20, F<sub>1,18</sub>=29806.11.25, P<0.0001 and n=20, F<sub>1,18</sub>=21925.4, P<0.0001, respectively). See text for statistical methods.

## DISCUSSION

Our results regarding the impact of esca on stem xylem integrity show that the presence of plant-derived tyloses induced hydraulic failure in 60% of symptomatic stems of the current year. Tyloses were only observed in symptomatic stems, and resulted in more than 50% PLC in 40% of the stems, unrelated to symptom age. We demonstrated that the presence of leaf symptoms during previous seasons had no impact on the likelihood of symptom appearance in the current year, or on stem hydraulic conductivity and xylem anatomy. Vascular fungi were never detected in the same organs as the tyloses (stems of the current year), and although they were present in trunks of both tiger-stripe and control plants, tiger-stripe plants showed higher quantities of fungal DNA. Among control plants that did not express symptoms in the year of the study, we found higher quantities of fungal DNA in trunks of those plants with a long-term history of symptom formation. Albeit xylem occlusions were not observed in the totality of tiger-stripe stems, they could amplify yield loss plant mortality, especially in the context of climate change as they impair water transport in a majority of symptomatic stems.

### *In vivo* xylem integrity observations and hydraulic vulnerability segmentation

Using direct X-ray microCT imaging in esca symptomatic stems, we found that hydraulic conductivity loss was almost entirely associated with the presence of tyloses. Different studies have investigated the link between vascular pathogen development and hydraulic conductivity in stems (Collins *et al*., 2009; Lachenbruch and Zhao, 2019; Mensah *et al*., 2020). During biotic stresses, air embolisms have been shown to decrease hydraulic conductivity during bacterial leaf scorch disease (McElrone *et al*., 2003; 2008), Pierce’s disease (Pérez-Donoso *et al*., 2016), and Pine wilt disease (Yazaki *et al*., 2018). In the case of fungal wilt diseases, the hydraulic conductivity loss was associated with nongaseous embolism (i.e. tyloses) at the point of pathogen inoculation (Guerard *et al*., 2000; Sallé *et al*., 2008; Beier *et al*., 2017; Mensah *et al*., 2020), or with canker presence in naturally infected stems (Lachenbruch and Zhao, 2019).

Using iohexol we were able to visually observe the exact spatial organization of functional vessels. Interestingly, in some symptomatic samples we found functional vessels surrounding the non-functional xylem (Fig. 2J-L), suggesting that the plant was able to preserve the more external vessels from occlusions or to form new functional vessels after the loss of conductivity. Moreover, the sectoriality of the occlusions observed in Fig. 2J-L was reminiscent of the sectoriality observed in the distributions of trunk necrosis, especially on the brown stripe necrosis appearing along the vasculature (Lecomte *et al*., 2012).

Comparing these results with our precedent study using the same technique in leaves, we showed that esca symptomatic leaves presented higher levels of occlusion PLC (61 ± 7% in midribs, and 54 ± 9% in petioles, data from Bortolami *et al*., 2019) compared to stems (27 ± 8 %, occlusion PLC), suggesting hydraulic vulnerability segmentation (although PLC in leaves and stems were measured in different plants and years). The hydraulic segmentation theory relies on the fact that annual organs (i.e. leaves) are more vulnerable than perennial organs (i.e. stems) to drought induced air embolism (Tyree and Ewers, 1991). Grapevine is well known for exhibiting strong hydraulic vulnerability segmentation (Charrier *et al*., 2016; Hochberg *et al*., 2016; 2017). This is thought to be adaptive, where the higher vulnerability in leaves and petioles favors embolism formation and leaf shedding prior to embolism formation in stems, thus protecting the perennial organs. Our observations during esca pathogenesis demonstrate that, analogous to the hydraulic vulnerability segmentation theory, leaves appear more vulnerable to the formation of nongaseous embolism as well, which could mitigate the risk of hydraulic failure in perennial organs. From another perspective, the difference may not be a direct effect of the specific organ’s vulnerability to nongaseous embolism, but a consequence of a difference in the accumulation of putative toxins and/or elicitors. Indeed, we confirmed here that esca leaf symptoms occur at a distance from the pathogen niche because vascular pathogens were never detected in stems of the current year, suggesting that the plant may transport a signal (i.e. toxins or elicitors) from the infected trunk up to the leaves. If the signal accumulates in leaves in a higher amount than it does in the stems (water potentials are more negative in leaves compared to stems), and stimulates occlusion formation, stems would then be secondarily affected.

### Hydraulic conductivity, tyloses, and vessel anatomy

Tyloses could have different impacts, both positive and negative, during wilt disease pathogenesis: (i) tyloses contribute to pathogen resistance as they aim to seal off vessel lumens and impede pathogens spread throughout the host (CODIT model, Shigo, 1984). This is the case regarding the susceptibility of different species or varieties to specific pathogens (Jacobi and MacDonald, 1980; Ouellette *et al*., 1999; Clérivet *et al*., 2000; Et-Touil *et al*., 2005; Venturas *et al*., 2014; Park and Juzwik 2014; Rioux *et al*., 2018), in particular to *Phaeomoniella chlamydospora*, one of the pathogen associated with esca (Pouzoulet *et al*., 2017; 2020). (ii) In other studies, it has been shown that tyloses can exacerbate symptoms (Talboys, 1972): they cause a reduction in stem hydraulic conductivity, sometimes associated with a reduction in stomatal conductance in leaves and, in the most severe cases, wilting (Parke *et al*., 2007; Beier *et al*., 2017; Lachenbruch and Zhao, 2019, Mensah *et al*., 2020 during fungi development; Sun *et al*., 2013; Deyett *et al*., 2019 during Pierce’s disease). Our results suggest that during esca tyloses might lead to symptom exacerbation. Esca has also been suggested to lead to a general reduction in xylem water transport and stomatal conductance (Ouadi *et al*., 2019), and tyloses could be a major contributor to these phenomena as during winter senescence (Salleo *et al*., 2002; Sun *et al*., 2008). However, when symptomatic stems have no tyloses (~37% of the stems with tiger-stripe symptoms), esca leaf symptom formation seems to arise from within the leaf itself, and may not result from upstream hydraulic failure. Although tyloses were never detected in asymptomatic stems prior to the onset of leaf symptoms, the time sequence of tylose and leaf symptom development has still to be determined. Since both the microCT and anatomical observations visualize relatively narrow regions of the stems, tylose presence could have been underestimated (i.e. if there was additional tylose development up or downstream of the stem sections visualized). However, it should be pointed out that if significant underestimation were present we would expect some loss of conductivity even in internode sections from which we observed no tyloses in the sampled cross sections. At least when considering a single internode our direct hydraulic conductivity measurements do not support the hypothesis that tyloses were underestimated (Fig. 4B).

Xylem is the battleground between vascular pathogens and the plant’s defense response (Yadeta and Thomma, 2013). Even if xylem vessel anatomy is less investigated, it could have a crucial role in plant resistance and response to vascular pathogens. For example, during Dutch elm wilt disease (due to *Ophiostoma spp*.) the most sensitive species and varieties present wider xylem vessels (Elgersma, 1970; Mcnabb *et al*., 1970; Solla and Gil 2002; Pita *et al*., 2018). Smaller vessels could occlude faster, sustaining a more efficient pathogen restriction (Venturas *et al*., 2014). Our results on xylem vessel anatomy suggest that stems with tyloses tend to present higher densities of small vessels, even if we did not observe any differences in total *k_th_* values and microCT scans showed that occlusions appear randomly in every vessel size class (Fig. S3). It could be possible that tylose formation might be interfering with stem water relations reducing the carbohydrates available for plant growth, producing smaller vessels in stems of symptomatic plants. In contrast, artificial inoculations showed that xylem vessel diameter had a strong impact on esca-related vascular pathogen development (Pouzoulet *et al*., 2017; 2020), and in the kinetic of vessel occlusion in grapevine stems (Pouzoulet *et al*., 2019). The relationships between esca leaf symptoms, xylem anatomy, and tylose presence should be studied in detail in trunks, where vascular pathogens are present, and among different grapevine varieties and rootstocks as they are known to show different susceptibility to symptom expression.

### Long-term consequences of esca on leaf symptom expression and stem hydraulic integrity

In field surveys, esca leaf symptoms are often randomly distributed spatially throughout vineyards and are not consistent from season to season in individual vines (Mugnai *et al*., 1999; Surico *et al*., 2000; Marchi *et al*., 2006; Guerin-Dubrana *et al*., 2013; Li *et al*., 2017). However, esca-related vine death is strongly related to leaf symptoms as death is usually observed following a year with symptom expression (Guerin-Dubrana *et al*., 2013). In agreement with these field studies, we observed similar percentages of symptomatic plants between those that had already expressed esca symptoms in the past (from one to seven consecutive years, pS plants), and those that had never expressed symptoms over the past seven years (pA plants). However, we also found that pS plants expressed symptoms earlier in the season than pA plants, suggesting that symptoms might require more time to develop in pA plants. We did not find any significant differences in *k_s_* and *k_th_* values between plants with contrasted long-term symptom history. This result suggests that esca leaf symptoms may have xylem anatomical consequences within the year of expression by the production of tyloses, but not across seasons. Moreover, we showed that DNA pathogen amount (*Phaeoacremonium minimum* and *Phaeomoniella chlamydospora*) depends on the symptom expression in the season of sampling, and on the long-term symptom history. Altogether, these results suggest that a higher amount of vascular fungi in the trunk represents a higher risk in reproducing leaf symptoms, and consequently, a higher risk of plant death.

### Hydraulic failure and esca leaf symptom pathogenesis

Our results showed that, even if esca-related stem occlusion was extremely variable, 40% of the microCT analyzed stems presented a total PLC greater than 50%. Under drought conditions alone, studies suggest that grapevines are not able to recover in the current season from PLC greater than 50% in stems (Charrier *et al*., 2018). Thus, to what extent these levels of esca-induced hydraulic failure compromise future vine performance, and/or increase the likelihood of developing esca leaf symptoms in the future remains an open question.

We showed that, similarly to visual leaf symptoms, tyloses in stems were generated at a distance from the pathogen niche in the trunk. Comparing our results with Bortolami *et al*. (2019), we show that the PLC due to the occlusions (hydraulic failure) observed using microCT in leaves was on average twice higher than the PLC observed in stems in the present work. We could hypothesize that, following pathogen activities in the trunk, a signal passing through the xylem network and stimulating tyloses, first accumulates in leaves and then affects the stems. However, the exact signal and action remain unknown, as we showed that the presence of tyloses depended upon given symptomatic stems rather than symptomatic plants (i.e. two stems in the same plant, with same tiger-stripe symptoms, sampled at the same moment, could or could not present tyloses).

We showed that there were no differences in symptom expression, nor in the stem hydraulic properties, regarding the long-term symptom history. We can conclude that the processes that generate tiger-stripe symptoms are largely restricted to the current year of the symptom expression. However, in plants expressing symptoms for the first time according to our disease record, these processes could require more time, as they showed symptoms only late in the season. The presence of occlusion, leading to hydraulic failure in stems, could exacerbate leaf symptom expression in the following seasons, possibly contributing to death. We could speculate that a stem expressing extensive hydraulic failure could be more prone to express symptoms in the following year or, in the worst cases, to die. If the level of hydraulic failure could affect the stem mortality in the following year, the choice of stems with a complete absence of failure during the winter pruning could reduce the impact of esca in vineyards. The pruning practices are known to impact the course of infection and leaf symptom development and it has been shown that trunk renewal could be an effective management practice to prevent grapevine trunk diseases in the vineyard (Travadon *et al*. 2016, Kaplan *et al*. 2016, Gramaje *et al*. 2018). In addition, the presence of occlusions could also amplify plant susceptibility to drought-induced hydraulic failure, enhancing the risk of plant mortality in the field as suggested by McDowell *et al*. (2008). It could be speculated that a decrease in soil water potential or a high evaporative demand, concomitant to esca-induced hydraulic failure, could embolize the remaining functional xylem vessels stopping the water flow and desiccating plant tissues (this could be the case in apoplectic plants for example). In perspective, future studies should investigate the link between pathogen activities and occlusion development, especially in trunks, and the subsequent hydraulic failure consequences on whole plant physiology.

## SUPPORTING INFORMATION

The following Supporting Information is available for this article:

**Fig. S1**. Two-dimensional reconstruction of longitudinal cross sections from X-ray microCT volumes of grapevine stems.

**Fig. S2**. Relationship between *k_s_* and *k_th_* in control plants.

**Fig. S3**. Vessel density and percentage of occluded vessels in tiger-stripe stems for different vessel diameter classes.

**Fig. S4**. Relationships between *k_s_* and *k_th_* in each stem symptom category.

**Table S1**. Effect of year of uprooting, internode analyzed, and sampling date on *k_s_* and *k_th_* in control stems.

**Table S2.** Calculated theoretical hydraulic conductivity (*k_th_* %), and hydraulic conductivity loss (PLC %) from X-ray microCT volumes of intact grapevine stems.

## DATA AVAILABILITY STATEMENT

Raw datasets are available in the INRAE dataverse: Bortolami, Giovanni; Farolfi, Elena; Badel, Eric; Burlett, Regis; Cochard, Herve; Ferrer, Nathalie; King, Andrew; Lamarque, Laurent J.; Lecomte, pascal; Marchesseau-Marchal, Marie; Pouzoulet, Jerome; Torres-Ruiz, Jose M.; Trueba, Santiago; Delzon, Sylvain; Gambetta, Gregory A.; Delmas, Chloe E.L., 2021, “Raw data for the paper “Seasonal and long-term consequences of esca on grapevine stem xylem integrity””, https://doi.org/10.15454/U9KJEW, Portail Data INRAE, V1.

## ACKNOWLEDGMENTS

We thank the experimental teams of UMR SAVE and UMR EGFV (Bord’O platform, INRAE, Bordeaux, France) and the SOLEIL synchrotron facility (beamline PSICHE) for providing the materials and logistics. Specifically, we thank Jérôme Jolivet and Sebastien Gambier (UMR SAVE) for providing technical knowledge and support for plant transplantation and maintenance. This work was supported by the French Ministry of Agriculture, Agrifood, and Forestry (FranceAgriMer and CNIV) within the PHYSIOPATH project (program Plan National Dépérissement du Vignoble, 22001150-1506) awarded to C.E.L.D., and program Investments for the Future (ANR-10-EQPX-16, XYLOFOREST).

## AUTHOR CONTRIBUTIONS

C.E.L.D., G.A.G., G.B., and S.D. designed the experiments;

E.F., G.A.G., S.D., E.B., R.B., H.C., A.K., L.J.L., J.M.T.-R., S.T. participated in synchrotron campaigns;

G.B., C.E.L.D., E.F., and N.F. conducted the esca symptom notations;

G.B., M.M.-M.,and N.F. conducted the histological observations;

E.F. conducted the hydraulic conductivity measurements and participated to data analyses;

N.F., and J.P., conducted the pathogen detection;

G.B. analyzed the microCT, optical images, and analyzed the data;

P.L. provided data on disease history of the plants;

G.B., C.E.L.D., and G.A.G. wrote the article;

all authors edited and agreed on the last version of the article

