## Supporting Information for "Seasonal and long-term consequences of esca on grapevine stem xylem integrity"

Giovanni Bortolami, Elena Farolfi, Eric Badel, Régis Burlett, Hervé Cochard, Nathalie Ferrer, Andrew King, Laurent J. Lamarque, Pascal Lecomte, Marie Marchesseau-Marchal, Jérôme Pouzoulet, José M. Torres-Ruiz, Santiago Trueba, Sylvain Delzon, Gregory A. Gambetta, Chloé E. L. Delmas

The following Supporting Information is available for this article:

**Fig. S1**. Two-dimensional reconstruction of longitudinal cross sections from X-ray microCT volumes of grapevine stems.

**Fig. S2**. Relationship between $k_s$ and $k_{th}$ in control plants.

**Fig. S3**. Vessel density and percentage of occluded vessels in tiger-stripe stems for different vessel diameter classes.

**Fig. S4**. Relationships between $k_s$ and $k_{th}$ in each stem symptom category.

**Table S1**. Effect of year of uprooting, internode analyzed, and sampling date on $k_s$ and $k_{th}$ in control stems.

**Table S2.** Calculated theoretical hydraulic conductivity ($k_{th}$ %), and hydraulic conductivity loss (PLC %) from X-ray microCT volumes of intact grapevine stems.

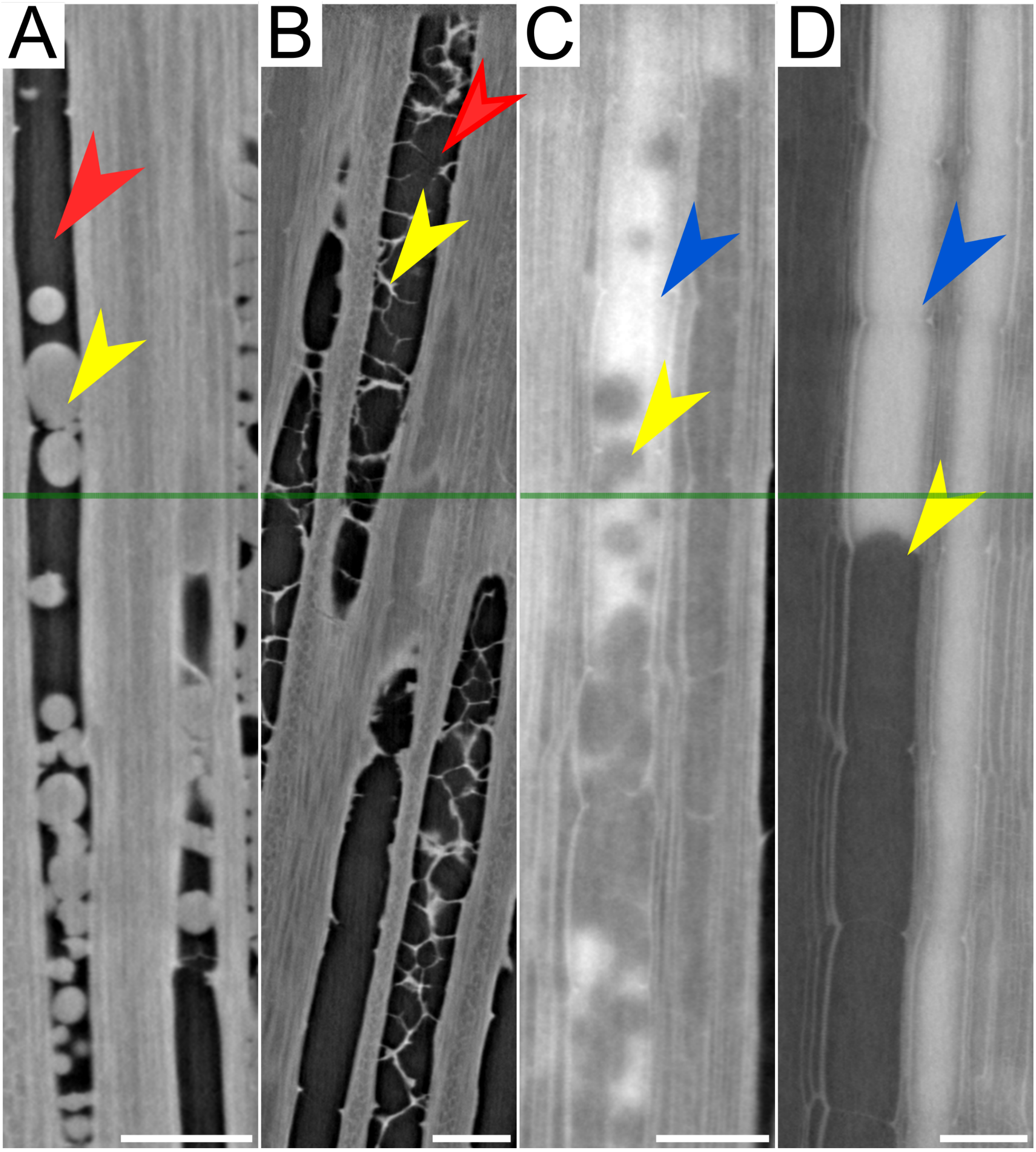


**Fig. S1**. Two-dimensional reconstruction of longitudinal cross sections from X-ray microCT volumes of grapevine stems, examples of apparently air-filled (A, B; red arrowheads), and apparently functional (C, D; blue arrowheads) vessels. Tyloses can only partially occlude the vessels (A, C, D, yellow arrowheads), or only the tylose walls are present without cellular content (B, yellow arrowheads). If only transversal cross sections (e.g. green lines) are analyzed, those vessels could be mistakenly considered as non-occluded. Scale bars = 200µm.

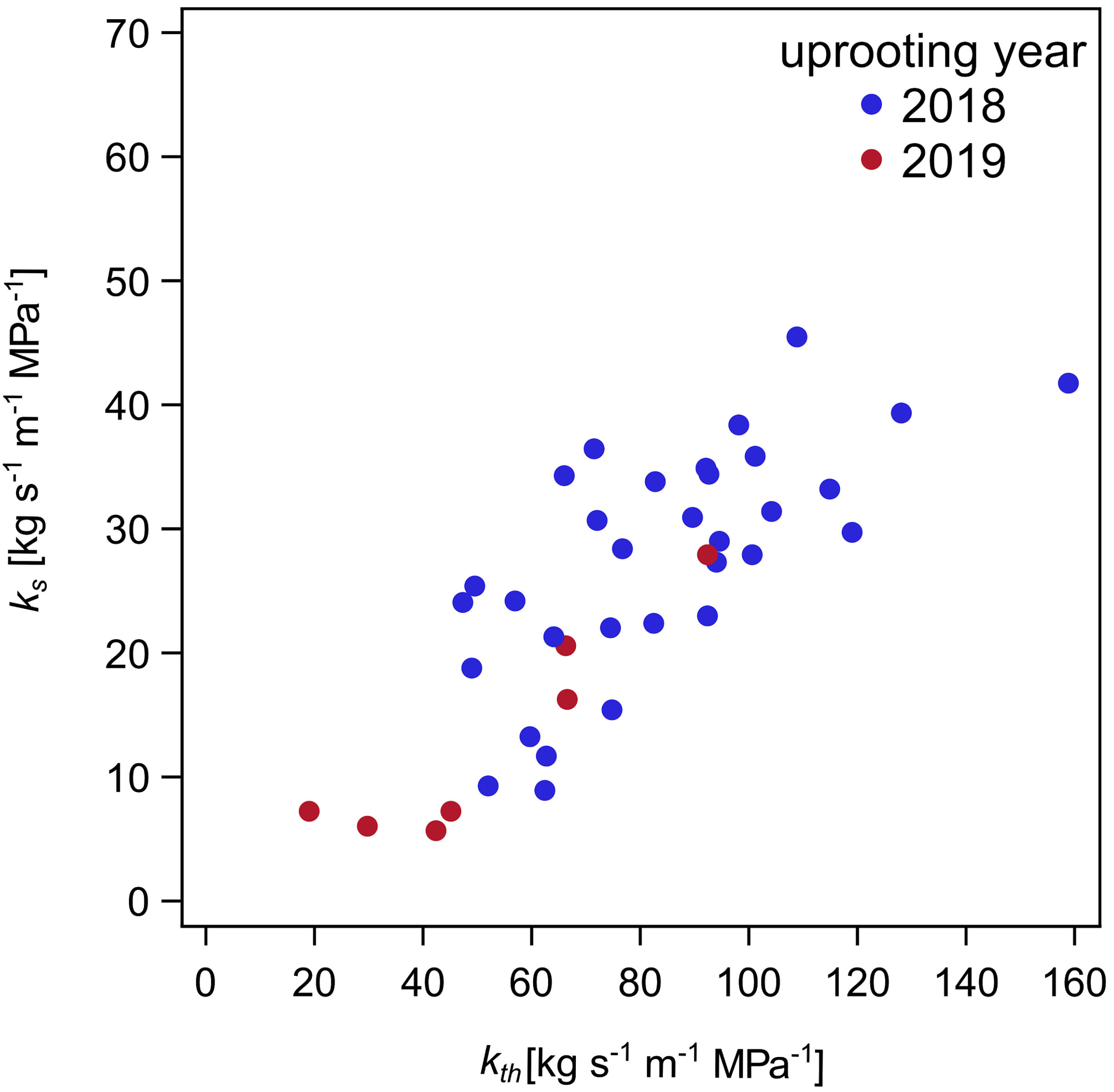


**Fig. S2**. Relationship between $k_s$ and $k_{th}$ in control plants. Blue dots represent plants uprooted in 2018 and red dots represent plants uprooted in 2019.

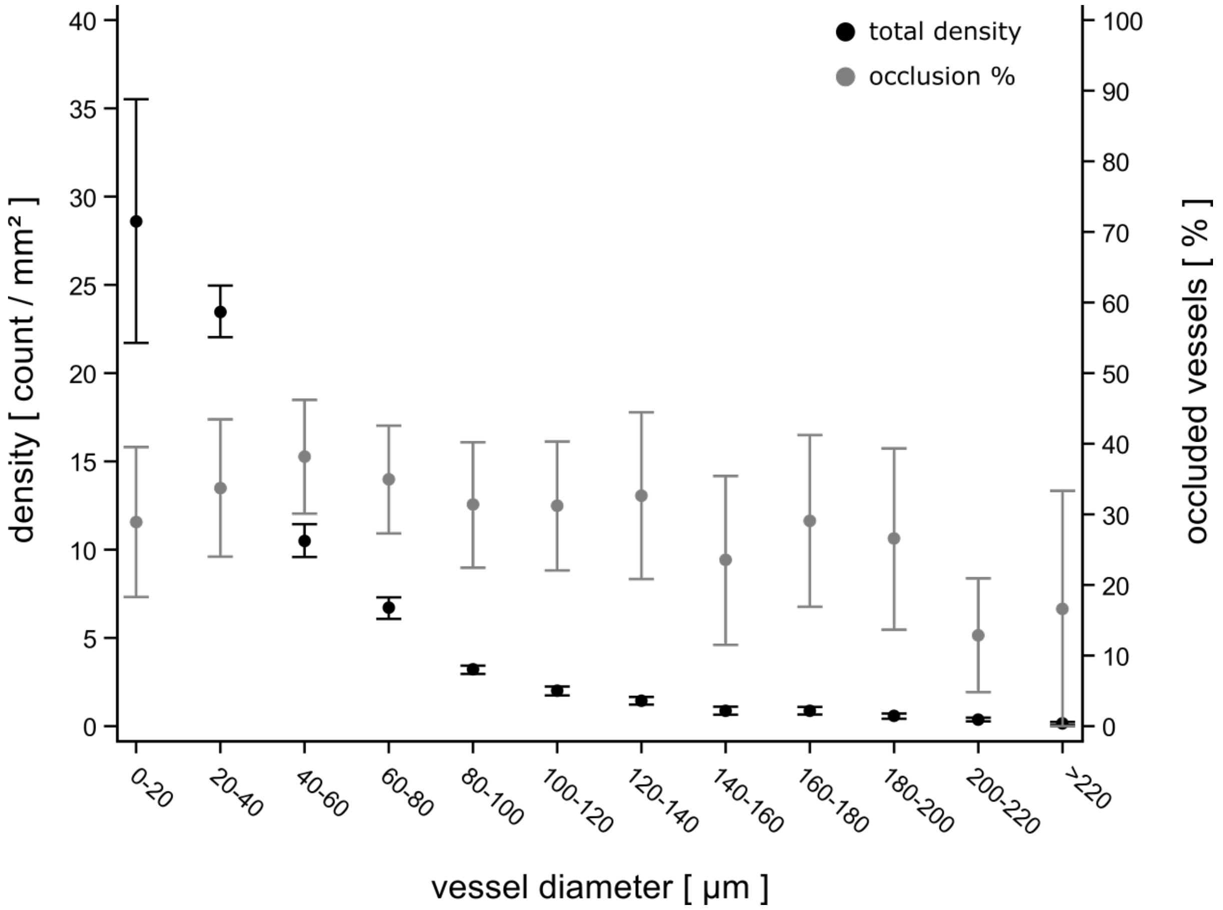


**Fig. S3**. Mean vessel density (black circles) in tiger-stripe stems for different vessel diameter classes and mean percentage of occluded vessels (grey circles) in each class from X-ray microCT imaging analysis. Error bars represent standard errors.

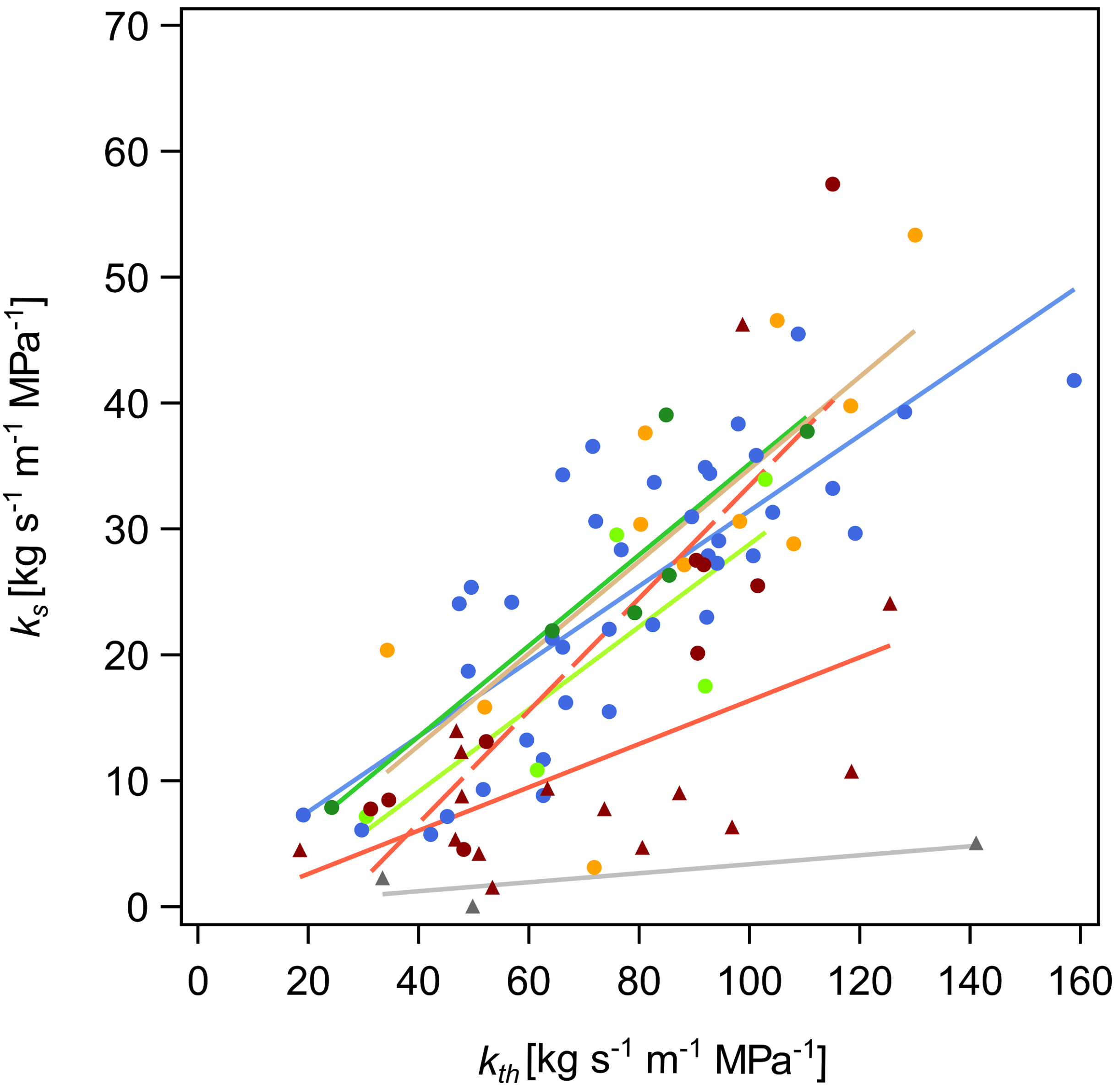


**Fig. S4**. Relationships between $k_s$ and $k_{th}$. Blue symbols represent measurements in control stems; dark green symbols represent asymptomatic stems in plants before symptom appearance; yellow symbols represent pre-symptomatic stems in plants before symptom appearance; light green symbols represent asymptomatic stems in plants after symptom appearance; red circles and dashed red line tiger-stripe stems without tyloses; red triangles and solid red line tiger-stripe stems with tyloses; grey symbols represent apoplectic stems. Sample sizes, $R^2$ and equations of regression lines are presented in Table 2.

**Table S1**. Effect of year of uprooting, internode analyzed, and sampling date on $k_s$ and $k_{th}$ in control stems (n=39 stems from 23 plants) .

| **Fixed effects** | $k_s$ **(n=39)** | $k_{th}$ **(n=39)** |
|---|---|---|
| Year of uprooting | $\mathbf{F_{1,16} = 10.31}$<br>$\boldsymbol{P} \mathbf{= 0.006}$ | $\mathbf{F_{1,16} = 9.38}$<br>$\boldsymbol{P} \mathbf{= 0.007}$ |
| Internode | $F_{4,12} = 2.15$<br>$P = 0.14$ | $F_{4,12} = 2.22$<br>$P = 0.13$ |
| Sampling date | $F_{7,9} = 1.77$<br>$P = 0.21$ | $F_{7,9} = 1.53$<br>$P = 0.27$ |

Individual generalized linear mixed models with the individual plants entered as a random effect in the models. Statistically significant results (P<0.05) are shown in bold.

**Table S2.** Calculated theoretical hydraulic conductivity ($k_{th}$ %), and hydraulic conductivity loss (PLC %) from X-ray microCT volumes of intact grapevine stems.

| Stems | n | Functional $k_{th}$ | Native PLC | Occlusion PLC |
|---|---|---|---|---|
| Control | 3 | 92.75 ± 2.60 | 6.54 ± 2.61 | 0.71 ± 0.02 |
| Esca (tiger-stripe) | 10 | 60.22 ± 9.70 | 12.25 ± 2.87 | 27.53 ± 8.24 |

Values are means ± standard error, n = sample size (stems)
